# Impact of Latent Tuberculosis on Inflammatory Biomarkers in Crohn’s Disease: A Comparative Study of Cytokines, CRP, and Calprotectin

**DOI:** 10.1101/2025.08.07.669228

**Authors:** Anmar Layth Talib, Jabbar S. hassan, Fadhil Abdullah Al Abbudi, Shaimaa M. Shehab, Marwah J. Kadhim

## Abstract

Latent tuberculosis (LTB) in Crohn’s disease patients could impact the inflammatory profile and treatment approach. Many cytokines, such as IL12, IL23, and TNF-α, are critical in both Crohn’s disease and mycobacterial infections; the exclusion of LTB-positive patients is important before initiation of immunotherapy because it may influence disease activity and inflammatory markers. This study compared and evaluated the levels of pro-inflammatory cytokines (TNF-α, IL-12, and IL-23) and fecal calprotectin and CRP in sick persons with Crohn’s disease who have latent TB infection and have not received immune-based therapy, and those who do not. The current research involved the enrollment of 100 patients with inflammatory bowel disease (IBD). Among them, 25 patients were diagnosed with latent tuberculosis infection (LTBI) based on the Gold-Gold-interferon-gamma release assay (IGRA). Patients who showed an IGRA negative (75), later received immunotherapy and were considered as the treated group. The Inflammatory markers (CRP and fecal calprotectin) and cytokine levels (TNF-α, IL-12, and IL-23) have been measured and compared between the treated group (n=75) and the untreated group (n=25). No significant correlation has been identified between the IGRA test and different cytokines. Smokers’ patients show higher levels of biomarker vs. non-smokers. Crohn’s disease patients who underwent treatment, including aminosalicylates (5-ASA), corticosteroids, and immunomodulators, had statistically significant differences in Inflammatory biomarker levels, calprotectin, interleukin-23 (IL-23), and interleukin-12 (IL-12), compared to those who did not receive any treatment, in which all biomarkers were markedly elevated in untreated patients. Fecal calprotectin levels showed statistically significant positive correlations with all measured inflammatory biomarkers. This study reported that Latent Tuberculosis infection in Crohn’s disease (CD) could not significantly affect the level of TNF-α, IL-12, IL-23, CRP, and fecal calprotectin. Form other hand, pro-inflammatory cytokines and calprotectin levels are significantly impacted by immune-based therapy.

## Introduction

Crohn’s disease (CD) is a chronic illness affecting the bowels system which occurs from the interaction and related with the genetic, immunological and environmental factors. One of the most important characteristics of this disease is disequilibrium in immune response, which induces excessive production of many proinflammatory cytokines, particularly interleukin-6 (IL-6), interleukin-1 beta (IL-1β), and tumor necrosis factor alpha (TNF-α), in addition to the vital role of these cytokine in the intestinal inflammation it’s also serve as a biomarker for disease activity [1].

Tuberculosis (TB) is an infectious disease which causes by Mycobacterium Tuberculosis and it’s a global problem issue, the progression of this disease is vary rang from acute primary tuberculosis to elimination of the pathogen and in many cases the host immune system can confined this bacterial agent and stop it from growing further by granuloma development and in these cases, it’s called dormant tuberculosis infection or latent (LTBI). This type of tuberculosis is asymptomatic, but the risk of reactivation and development of active tuberculosis remains high, particularly in immunosuppressed individuals [2].

A significant element of Crohn’s disease treatment is immunomodulation, which focuses on important inflammatory pathways to regulate disease activity in addition to targeted cytokine inhibition provided by biologic agents such as anti-TNF medications to prevent remission and avoid recurrence [3].

Moreover, TB infection has the ability to alter host immunological responses, which may have an impact on CD pathogenesis and cytokine expression. Despite being crucial for infection management, anti-TB treatment may also have an effect on the inflammatory environment, which could influence CD patients’ disease activity [4].

Due to the clinically significant interactions between immunological factors, TB infection, anti-TB medication, and CD disease activity, a wide range of studies is being conducted to understand this dilemma [5].

The aim of this study is to evaluate and compare the levels of fecal calprotectin, CRP, and pro-inflammatory cytokines (TNF-α, IL-12, and IL-23) in patients with Crohn’s disease who have latent Tuberculosis infection and those who do not.

## Methodology

### Study design

This is an observational, cross-sectional study of Inflammatory bowel illness patients who were referred for interferon gamma releasing assay (IGRA) testing to exclude latent tuberculosis (LTB), before initiating treatment with immune-based therapy (corticosteroid, anti-TNF, and mesalamine). This study was conducted from May 22, 2025, to July 25, 2025, at four academic Teaching Hospitals in Baghdad, Iraq. During this period, 100 IBD patients were enrolled in this study; of these, 25 patients were diagnosed as LTBI using by Gold IGRA (TB1 and TB2) with a control negative (nil) tube and a control positive (mitogen) tube. To evaluate the features of IBD patients, a retrospective review was carried out, and the clinical characteristics and results of the rate of LTB are documented and contrasted with non-LTB.

The interferon gamma releasing assay (IGRA) has been performed in accordance with the manufacturer’s guidelines (QIAGEN, Germany). Additionally, Cytokine levels (IL-12, IL-23, and TNF-α) and inflammatory markers (CRP and fecal calprotectin) were measured in all patients using an ELISA assay according to the manufacturer’s protocol.

The College of Medicine at Al-Nahrain University’s Institutional Review Board granted approval for this study on May 21, 2025, with number 20250591. Prior to participation, each adult participant gave their informed consent. After being fully informed about the study’s objectives, methods, and confidentiality, verbal informed consent was acquired. As authorized by the ethics committee, the study team recorded the permission and carried out the procedure in front of a witness.

### Data collection

Baseline demographic and clinical parameters were assessed, including smoking status, age at IBD diagnosis, gender, and underlying comorbidities like diabetes mellitus, hypertension, cancer, and pulmonary illness. The following extra IBD-related data were gathered: IBD kind and duration; anti-TNF therapy type and duration, mesalamine and other corticosteroids within the 3 months of sample collection.

### Inclusion Criteria

- Patients were diagnosed with Crohn’s disease using clinical, endoscopic, and histopathological criteria.
- No prior exposure to immunomodulating drugs (anti-TNF, corticosteroids, mesalamine)
- Patients who underwent IGRA testing

### Exclusion Criteria

- Patients with active TB at baseline
- Individuals with known autoimmune diseases other than Crohn’s
- Patients on immunosuppressive therapy that could influence cytokine levels
- Incomplete data on IGRA results

### Statistical analysis

The Statistical Package for the Social Sciences (SPSS) version 20 was used to analyze all of the data. Binomial data were presented as frequencies and percentages. To analyze proportional differences, the Chi-square (χ^2^) test was used. P-values below 0.05 were regarded as statistically significant.

## Results

### Descriptive Statistics and IGRA Results

Out of 100 Crohn’s patients enrolled in this study, 54 were Male, 46 were Female, with a male-to-female ratio of 1.17:1. According to smoking history, 58 (58%) were smokers and 42(42%) as non-smokers

Among those patients, 25 %(n=25) tested positive for latent tuberculosis by IGRA test with a result greater than 0.35 IU/mL. The remaining 75% (n = 75) had IGRA results < 0.35 IU/mL, suggesting no evidence of latent infection (Fig 1).

**Fig 1.**
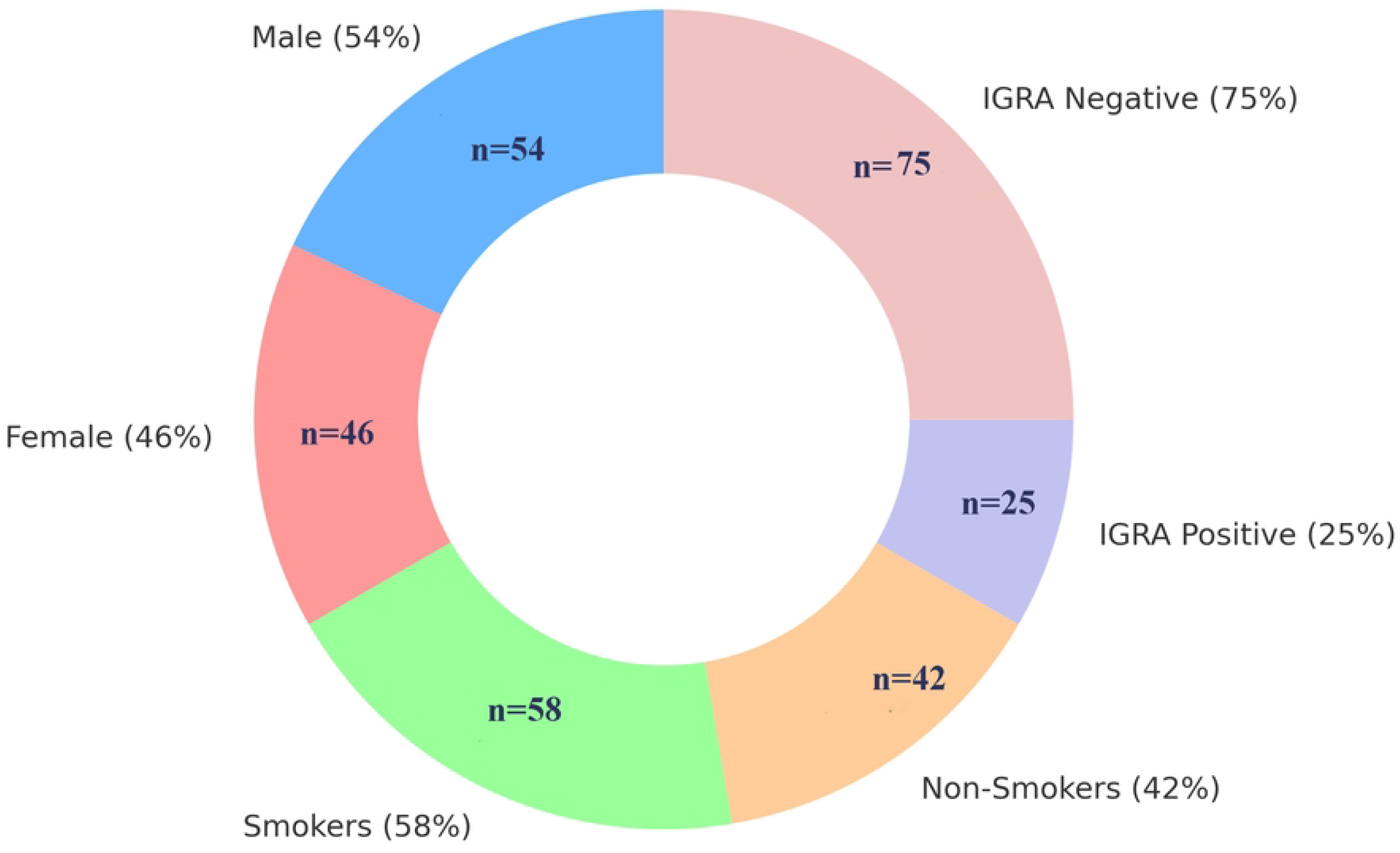
Patient Characteristics and Gold IGRA Test Results.

Cytokine levels (IL-12, IL-23, TNF-α) and inflammatory markers (CRP and fecal calprotectin) were compared between 75 patients who showed IGRA negative and received immunotherapy and the untreated group (n=25). In addition to that smoking impact was evaluated. The outcomes are proposed as mean ± standard deviation (SD) with p-values from statistical comparisons. There is a significant reduction in all biomarkers between the untreated and treated groups (*p < 0*.*05*). Smokers show higher levels of biomarker vs. non-smokers (*p < 0*.*05*) (Table 1).

**Table 1:** Biomarker Levels and Smoking by Treatment Status.

| Biomarker | Untreated<br>(Mean ±<br>SD) | Treated<br>(Mean ±<br>SD) | p-value | Smokers<br>(Mean ±<br>SD) | Non-Smokers<br>(Mean ± SD) | p-value<br>(Smoking) |
| --- | --- | --- | --- | --- | --- | --- |
| CRP (mg/dl) | 18.45 ±<br>14.32 | 13.27 ±<br>10.56 | 0.028 | 17.21 ±<br>12.87 | 12.83 ± 10.95 | 0.021 |
| TNF-α<br>(pg/mL) | 12.84 ±<br>8.91 | 8.56 ± 5.23 | 0.002 | 11.92 ±<br>7.84 | 7.65 ± 5.12 | <0.001 |
| **IL-12**<br>**(pg/mL)** | 11.73 ±<br>9.45 | 6.82 ± 5.67 | 0.001 | 10.85 ±<br>8.76 | 6.24 ± 4.93 | 0.003 |
| **IL-23**<br>**(pg/mL)** | 30.15 ±<br>11.24 | 22.47 ± 9.86 | 0.003 | 28.43 ±<br>10.67 | 21.15 ± 8.72 | <0.001 |
| **Calprotectin**<br>**(µg/g)** | 281.6 ±<br>198.4 | 148.3 ±<br>125.7 | <0.001 | 264.8 ±<br>187.2 | 132.5 ± 114.9 | <0.001 |

Regarding the correlation between the IGRA test and different cytokines included in this study, no significant correlations were observed between IGRA results (TB1 and TB2) and levels of CRP, fecal calprotectin, TNF-α, IL-12, or IL-23 in Crohn’s disease patients. All *p*-values exceeded 0.05 (Table 2).

**Table 2:** Correlation Between IGRA Test Results and Inflammatory Biomarkers.

| **Inflammatory Biomarker** | **Gold IGRA TB1 IU/mL**<br>**(<0.35)** | **Gold IGRA TB2 IU/mL**<br>**(<0.35)** |
| --- | --- | --- |
| **C-Reactive Protein mg/dl (<6.0)** | **r = 0.110, p = 0.277** | **r = 0.127, p = 0.210** |
| **Calprotectin titer in stool µg/g**<br>**(<43.2)** | **r = -0.018, p = 0.859** | **r = -0.029, p = 0.776** |
| **TNF-α (<8.1 pg/mL)** | **r = -0.014, p = 0.892** | **r = -0.022, p = 0.831** |
| **IL-12 (p70) (<5.0 pg/mL)** | **r = 0.097, p = 0.336** | **r = 0.064, p = 0.528** |
| **Interleukin-23 (<20 pg/mL)** | **r = -0.031, p = 0.756** | **r = -0.062, p = 0.540** |

r = correlation coefficient, p. *P*. value

Fecal calprotectin levels showed statistically significant positive correlations with all measured inflammatory biomarkers, which suggests associations between mucosal inflammation and systemic inflammatory responses in Crohn’s disease patients (Table 3).

**Table 3:** Calprotectin and pro-inflammatory cytokine levels (TNF-α, IL-12,IL-23, CRP) in Crohn’s disease patients.

| **Inflammatory Biomarker** | **Calprotectin titer in stool µg/g (<43.2)** |  |
| --- | --- | --- |
|  | **Correlation**<br>**coefficient** | ***P* value** |
| **C-Reactive Protein mg/dl (<6.0)** | 405** | *P* = 0.000 |
| **TNF-α (<8.1 pg/mL)** | 793** | *P* = 0.000 |
| **IL-12 (p70) (<5.0 pg/mL)** | 619** | *P* = 0.000 |
| **Interleukin-23 (<20 pg/mL)** | 673** | *P* = 0.000 |

Crohn’s disease patients that underwent treatment including aminosalicylates (5-ASA), corticosteroids, immunomodulators have statistically significant differences in Inflammatory biomarker levels calprotectin, interleukin-23 (IL-23), interleukin-12 (IL-12), TNF-α, compared to those that did not receive any treatment in which all biomarkers were markedly elevated in untreated patients (p < 0.001 for all comparisons) (Table 4).

**Table 4:** Serum Inflammatory and Fecal Biomarkers in Treated and Untreated Crohn’s Disease Patients.

| **Biomarkers** | **5-ASA** | **Corticosteroids** | **Immunomodulators** | **Untreated** | **p-value** |
| --- | --- | --- | --- | --- | --- |
| **Calprotectin**<br>**($\mu\text{g/g}$)** | 105.3<br>(88.1–<br>153.8) | 86.75 (60.6–119.4) | 109.1 (88.4–151.2) | 325.2 (311.7–<br>413.3) | <0.001 |
| **Interleukin-**<br>**23 (pg/mL)** | 22.87<br>(16.57–<br>27.75) | 18.9 (13.37–22.85) | 23.03 (16.12–26.05) | 34.32 (30.97–<br>38.85) | <0.001 |
| **IL-12**<br>**(pg/mL)** | 5.87<br>(4.74–<br>6.63) | 3.75 (2.83–5.94) | 5.99 (3.41–6.69) | 16.11 (13.31–<br>21.9) | <0.001 |
| **TNF-α**<br>**(pg/mL)** | 6.84<br>(5.06–<br>7.73) | 5.72 (4.42–6.96) | 5.95 (5.17–7.08) | 16.82 (13.86–<br>18.84) | <0.001 |

## Discussion

The cytokine responses in the inflammatory bowel diseases (IBDs) are considered as ample fruit that guides the highlighting of the mechanisms of these diseases; on the other hand, latent tuberculosis infection in Crohn’s disease patients potentially impacts the level of cytokine profile, including tumor necrosis factor alpha, interleukin-23, and interleukin-12, in addition to inflammatory markers like CRP and fecal calprotectin [6].

Moreover, immune-based therapy is one of the efficient strategies in the treatment of inflammatory bowel diseases (IBDs), but unfortunately, it’s related to LTBI reactivation and subsequently converts to active tuberculosis [7]. In this study, pro-inflammatory cytokine levels (TNF-α, IL-12, IL-23), CRP, and fecal calprotectin were evaluated and compared in patients with Crohn’s disease, with a focus on the presence of latent tuberculosis infection (LTBI), as determined by the interferon-gamma release assay (IGRA). The findings in this study may give a snapshot into both the role of proinflammatory mediators in the immunopathology of CD and the significance of LTBI in this population. This study was conducted with some limitations in that the treatment types, dose, or duration may affect the study outcome. Furthermore, this study is case case-control observational study which cannot establish causality (LTB causing cytokine changes vs. Crohn’s severity).

In the current study, it was found that among 100 Crohn’s patients, males were slightly predominant regarding females, with an around 1.17:1 male-to-female ratio. This result comes in accordance with those obtained in other studies in Asian and Middle Eastern populations [8,9].

While some western studies often show a more balanced sex ratio or a slight female predominance, which may be reflecting regional differences in genetic susceptibility or healthcare access [10,11]. Smoking in Crohn’s patients has a bad impact, which may be considered a factor that enhances disease severity and systemic inflammation and contributes to poor response to therapy. In this study, 58% of participants were former smokers. The same smoking finding was reported in other studies [12,13].

On the other hand, interferon-gamma release assay (IGRA) test revealed that 25% of patients tested positive for latent tuberculosis infection (LTBI), this percentage comes in agreement with previous studies in the Middle East, where LTBI prevalence among IBD patients ranges from 12% to 34%, [14], while the results of our study are higher than the result reported by Alayed et al [15], who mentioned that LTBI rate among IBD patients below 10%. Interferon-gamma release assay result findings are generally consistent with those reported in the above studies, but differences in sample size should be considered.

The study’s findings may highlight how important the IGRA exam and as a test that must be initiated in Crohn’s disease patients before starting immunotherapy. These demographic results, such as male predominance, high smoking percentage, and notable LTBI rates, may attract attention to the population with specific risk profiles that may influence both disease progression and therapeutic decision-making in Crohn’s disease management.

Concerning the comparison of CD patients’ biomarkers between the treated and untreated groups, according to this outcome, there was additionally a significant decrease in these biomarkers in CD patients undergoing immune-based therapy (aminosalicylates, corticosteroids, or immunomodulators) versus those patients who have not received such therapy. Many researchers state that immune-based therapy showed anti-inflammatory effects that led to a decrease in the level of TNF-α and the IL-12/23 cytokine axis; such results align with our findings [16-18].

Our data also demonstrated the impact of smoking on inflammatory biomarkers in CD patients, with significantly higher levels of TNF-α, IL-12, IL-23, CRP, and calprotectin than in non-smokers patients. AlQasrawi [19] reported such a finding, and they explained such result to the effect of smoking in the mucosa of intestinal tissue, which enhances mucosal inflammation. While Boronat et al. [20] attributed such a result to the effect of tobacco in upregulation of the TNF-α, IL-6, and IL-1β pathways, potentially exacerbating disease severity and resistance to standard therapies.

Although IGRA-negative people were the exclusive focus in this investigation, it is important to note that this population is deemed appropriate for the commencement of immunosuppressive medication. IGRA-negative individuals have a low risk of TB reactivation, according to Campainha et al [21] and Viladrich et al[22], confirming the safety of immunotherapy in this subgroup. Therefore, biomarker alteration in these results is due to immunologically based therapy rather than being confused by TB-related immunological activation.

When combined, these findings highlight how crucial biomarker monitoring is for directing treatment choices and forecasting CD outcomes. They also highlight the harmful effects of smoking, which can worsen inflammation and reduce the effectiveness of treatment, highlighting the therapeutic necessity of smoking cessation counseling in this population.

The study’s findings that fecal calprotectin levels and all tested inflammatory biomarkers, such as TNF-α, IL-12, IL-23, and C-reactive protein (CRP), are significantly correlated are particularly interesting. Indeed, fecal calprotectin is considered a non-invasive biomarker that in CD patients can be used to demonstrate mucosal inflammatory activity. It is worth noting that the strong correlation between these inflammatory biomarkers and Crohn’s disease development, particularly TNF-α, aids in the interlinking of mucosal inflammation. Correspondingly, many studies reported that elevated fecal calprotectin levels can be decline when using therapy that targets TNF-α. [23-25].

It is important to note that patients who received immunological therapy and those who did not showed significant differences in all biomarker levels under study. The levels of fecal calprotectin (median 325.2 µg/g), TNF-α (16.82 pg/mL), IL-23 (34.32 pg/mL), and IL-12 (16.11 pg/mL) were significantly higher in those not receiving immunological-based therapy, with p values < 0.001 in every comparison. In the same way, Van den Berghe et al [26] and Kumar et al [27] reported that biomarker-level differences exist between treated responder’s vs untreated or non-responders.

## Conclusion

According to this study, TNF-α, IL-12, IL-23, CRP, and fecal calprotectin levels were not significantly affected by latent tuberculosis infection in Crohn’s disease (CD). This may highlight that LTBI may not directly alter the inflammatory profile in CD patients. Form other hand, pro-inflammatory cytokines and calprotectin levels are significantly impacted by immune-based therapy, highlighting its efficacy in reducing mucosal and systemic inflammation in CD. The necessity for focused smoking cessation strategies in CD therapy, however, this study suggested that smoking is a modifiable risk factor that is strongly linked to higher biomarker levels and possibly worse disease outcomes.

## Acknowledgments

The authors would like to express deep appreciation to the Gastrointestinal (GIT) Medical Teaching Hospital in Baghdad, Iraq, for their invaluable support throughout this study. The collaboration, access to clinical data, and the dedicated efforts of the hospital’s medical staff significantly contributed to the successful completion of this work. We are especially thankful for the guidance and facilities provided, which created a conducive environment for academic research and scientific advancement.

## References

1. Zhao Y, Xu M, Chen L, Liu Z, Sun X. Levels of TB-IGRA may help to differentiate between intestinal tuberculosis and Crohn’s disease in patients with positive results. Therapeutic advances in gastroenterology. 2020 May;13:1756284820922003.

2. Sachdeva K, Kumar P, Kante B, Vuyyuru SK, Mohta S, Ranjan MK, et al. Interferon-gamma release assay has poor diagnostic accuracy in differentiating intestinal tuberculosis from Crohn’s disease in tuberculosis endemic areas. Intest Res. 2023 Apr 1;21(2):226–34.

3. Fehily SR, Al‐Ani AH, Abdelmalak J, Rentch C, Zhang E, Denholm JT, et al. latent tuberculosis in patients with inflammatory bowel diseases receiving immunosuppression— risks screening, diagnosis, and management. Alimentary pharmacology & therapeutics. 2022 Jul;56(1):6–27.

4. Kedia, S., Mouli, V.P., Kamat, N., Sankar, J., Ananthakrishnan, A., Makharia, G. et al. Risk of tuberculosis in patients with inflammatory bowel disease on infliximab or adalimumab is dependent on the local disease burden of tuberculosis: a systematic review and meta-analysis. Official journal of the American College of Gastroenterology| 2020, 115(3), pp.340–349.

5. Lee JY, Oh K, Hong HS, Kim K, Hong SW, Park JH, et al. Risk and characteristics of tuberculosis after anti-tumor necrosis factor therapy for inflammatory bowel disease: a hospital-based cohort study from Korea. BMC Gastroenterology. 2021 Dec; 21:1–9.

6. Friedrich M, Pohin M, Powrie F. Cytokine networks in the pathophysiology of inflammatory bowel disease. Immunity. 2019 Apr 16;50(4):992–1006.

7. Fehily SR, Al‐Ani AH, Abdelmalak J, Rentch C, Zhang E, Denholm JT, et al. latent tuberculosis in patients with inflammatory bowel diseases receiving immunosuppression— risks screening, diagnosis and management. Alimentary pharmacology & therapeutics. 2022 Jul;56(1):6–27.

8. Lee JW, Eun CS. Inflammatory bowel disease in Korea: epidemiology and pathophysiology. The Korean Journal of Internal Medicine. 2022 Jul 29;37(5):885.

9. Ng SC, Shi HY, Hamidi N, Underwood FE, Tang W, Benchimol EI, et al. Worldwide incidence and prevalence of inflammatory bowel disease in the 21st century: a systematic review of population-based studies. The Lancet. 2017 Dec 23;390(10114):2769–78.

10. Lungaro L, Costanzini A, Manza F, Barbalinardo M, Gentili D, Guarino M, et al. Impact of female gender in inflammatory bowel diseases: a narrative review. Journal of personalized medicine. 2023 Jan 17;13(2):165.

11. Goodman WA, Erkkila IP, Pizarro TT. Sex matters: impact on pathogenesis, presentation and treatment of inflammatory bowel disease. Nature reviews Gastroenterology & hepatology. 2020 Dec;17(12):740–54.

12. Chen BC, Weng MT, Chang CH, Huang LY, Wei SC. Effect of smoking on the development and outcomes of inflammatory bowel disease in Taiwan: a hospital-based cohort study. Scientific Reports. 2022 May 10;12(1):7665.

13. Wang P, Hu J, Ghadermarzi S, Raza A, O’Connell D, Xiao A, et al. Smoking and inflammatory bowel disease: a comparison of China, India, and the USA. Digestive diseases and sciences. 2018 Oct;63:2703–13.

14. Alhalabi M, Alshiekh HA, Alsaiad S, Zarzar M. Prevalence of opportunistic infections in Syrian inflammatory bowel disease patients on biologic therapy: a multi-center retrospective cross-sectional study. BMC Infectious Diseases. 2025 May 4;25(1):652.

15. Alayed M, Alrasheed A, Alenezi M, Binsalmah I, Almutairdi A, et al. Prevalence and Outcomes of Active and Latent Tuberculosis Among Patients with Inflammatory Bowel Disease (IBD) in an Endemic Region. Official journal of the American College of Gastroenterology| ACG. 2024 Oct 1;119(10S):S1084–5.

16. Verstockt B, Salas A, Sands BE, Abraham C, Leibovitzh H, Neurath MF. IL-12 and IL-23 pathway inhibition in inflammatory bowel disease. Nature Reviews Gastroenterology & Hepatology. 2023 Jul;20(7):433–46.

17. Aggeletopoulou I, Assimakopoulos SF, Konstantakis C, Triantos C. Interleukin 12/interleukin 23 pathway: Biological basis and therapeutic effect in patients with Crohn’s disease. World journal of gastroenterology. 2018 Sep 28;24(36):4093.

18. Sewell GW, Kaser A. Interleukin-23 in the pathogenesis of inflammatory bowel disease and implications for therapeutic intervention. Journal of Crohn’s and Colitis. 2022 Apr 1;16(Supplement_2):ii3–19.

19. AlQasrawi D, Qasem A, Naser SA. Divergent effect of cigarette smoke on innate immunity in inflammatory bowel disease: A nicotine-infection interaction. International Journal of Molecular Sciences. 2020 Aug 13;21(16):5801.

20. Boronat-Toscano A, Vañó I, Monfort-Ferré D, Menacho M, Valldosera G, Caro A, et al. Smoking suppresses the therapeutic potential of adipose stem cells in Crohn’s disease patients through epigenetic changes. Cells. 2023 Mar 27;12(7):1021.

21. Campainha S, Gomes T, Carvalho A, Duarte R. Negative predictive value of TST and IGRA in anti-TNF treated patients. European Respiratory Journal. 2012 Aug 31;40(3):790–1.

22. Viladrich IM, Tello ED, Solano-Lopez G. Consensus document on prevention and treatment of tuberculosis in patients for biological treatment. Archivos de Bronconeumología (English Edition). 2016 Jan 1;52(1):36–45.

23. Lucaciu LA, Ilieş M, Vesa ŞC, Seicean R, Din S, Iuga CA, Seicean A. Serum interleukin (IL)-23 and IL-17 profile in inflammatory bowel disease (IBD) patients could differentiate between severe and non-severe disease. Journal of personalized medicine. 2021 Nov 2;11(11):1130

24. Bourgonje AR, Martels JZ, Faber KN, Dijkstra G. Increased fecal calprotectin levels in Crohn’s disease correlate with elevated serum Th1-and Th17-associated cytokines. PLoS One. 2018 Feb 21;13(2):e0193202.

25. Langhorst, J., Elsenbruch, S., Koelzer, J., Rueffer, A., Michalsen, A. and Dobos, G.J., 2008. Noninvasive markers in the assessment of intestinal inflammation in inflammatory bowel diseases: performance of fecal lactoferrin, calprotectin, and PMN-elastase, CRP, and clinical indices. Official journal of the American College of Gastroenterology| ACG, 103(1), pp.162–169.

26. Van den Berghe, N., Alsoud, D., Verstockt, B., Vermeire, S., Declerck, P. and Thomas, D., 2023. Evaluation of serum cytokines and acute phase proteins as possible pharmacodynamic biomarkers to monitor endoscopic remission during ustekinumab therapy in patients with Crohn’s disease. Therapeutic Advances in Gastroenterology, 16, p.17562848231189110

27. Kumar M, Murugesan S, Ibrahim N, Elawad M, Al Khodor S. Predictive biomarkers for anti-TNF alpha therapy in IBD patients. Journal of Translational Medicine. 2024 Mar 16;22(1):284.

